# The plastid-nucleus located DNA/RNA binding protein WHIRLY1 regulates microRNA-levels during stress

**DOI:** 10.1101/197202

**Authors:** Aleksandra Swida-Barteczka, Anja Krieger-Liszkay, Wolfgang Bilger, Ulrike Voigt, Götz Hensel, Zofia Szweykowska-Kulinska, Karin Krupinska

## Abstract

In this article a novel mechanism of retrograde signaling by chloroplasts during stress is described. This mechanism involves the DNA/RNA binding protein WHIRLY1 as a regulator of microRNA levels. By virtue of its dual localization in chloroplasts and the nucleus of the same cell, WHIRLY1 was proposed as an excellent candidate coordinator of chloroplast function and nuclear gene expression (Grabowski et al., 2008; Foyer et al., 2014). In this study the putative involvement of WHIRLY1 in stress dependent retrograde signaling was investigated by comparison of barley (*Hordeum vulgare* L., cv. Golden Promise) wild-type and transgenic plants with an RNAi-mediated knockdown of *WHIRLY1*. In contrast to the wild type, the transgenic plants were unable to cope with continuous high light conditions. They were impaired in production of several microRNAs mediating post-transcriptional responses during stress (Kruszka et al., 2012, Sunkar et al., 2012). The results support a central role of WHIRLY1 in retrograde signaling and underpin a so far underestimated role of microRNAs in this process.

---

WHIRLY1 belongs to a small plant specific family of DNA/RNA-binding proteins. By immunological methods, WHIRLY1 has been detected in chloroplasts and the nucleus of the same cell (Grabowski et al., 2008). Accordingly, functions of WHIRLY1 were reported for both compartments. In chloroplasts of barley, WHIRLY1 was shown to be the major compacting protein of nucleoids (Krupinska et al., 2014). Moreover, WHIRLY1 has been found to bind to plastid RNAs (Melonek et al., 2010; Prikryl et al., 2008). In chloroplasts of *Arabidopsis thaliana*, WHIRLY1 was reported to maintain plastid genome stability (Maréchal et al., 2009). In the nucleus, WHIRLY1 was originally detected as a component of a transcriptional activator of the *PR10a* gene of potato (Desveaux et al., 2000). Furthermore, it has been found to bind to telomeres (Yoo et al., 2007).

Chloroplasts act as sensors of the environmental situation and produce diverse signals informing about the functionality of the photosynthetic apparatus (Pfalz et al., 2012, Kleine and Leister, 2016). These retrograde signals comprise redox changes and reactive oxygen species and regulate gene expression in the nucleus in particular during stress situations (Dietz, 2015). Although in recent years several compounds involved in chloroplast-to-nucleus communication have been identified, the full repertoire of molecular mechanisms adjusting nuclear gene expression to environmental cues remains obscure (Chan et al., 2016).

To investigate the impact of WHIRLY1 on stress resistance of barley plants, seedlings of three independent transgenic lines with an RNAi-mediated knockdown of *WHIRLY1* (RNAiW1-1, RNAi-W1-7 and RNAi-W1-9) were grown in continuous light at four different irradiances (50, 120, 200, 350 µmol photons m^-2^s^-1^). Leaves had reduced levels of the WHIRLY1 protein ranging from undetectable traces (RNAi-W1-7) to 10% of the wild-type level (RNAi-W1-1, RNAi-W1-9) (Krupinska et al., 2014). The reduction in length was the same in both lines having 10% the WHILRLY1. Therefore, only the results obtained for line W1-1 besides line W1-7 are presented in Figure 1. The reduction in leaf length occurred irrespective of the irradiance (Fig. 1A) indicating that WHIRLY1 has a general positive effect on growth.

**Figure 1.**
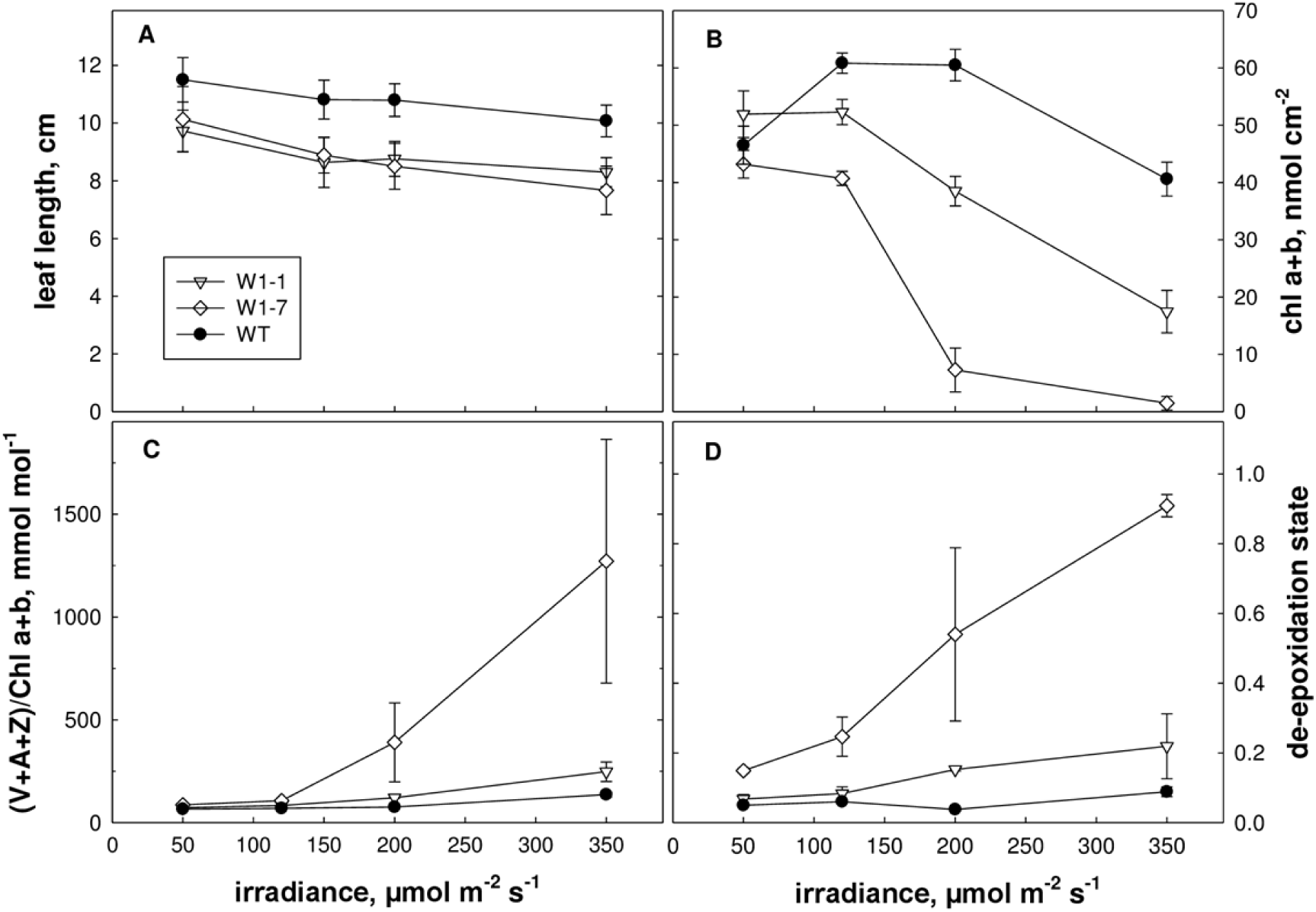
Characterization of *WHIRLYI* knockdown lines at the seedling stage. Seedling were exposed to continuous irradiation at 50, 120,200 or 350 µmol photons m^-2^ s^-1^ for 7 days. Lengths of the primary leaves (cm) are indicated (A). Pigment extracts from the wild type (WT) and the RNAi-W 1 lines (W-1,W-7 were compared by HPLC for the content of chlorophylls/leaf area(B), the radio of xanthophyll cycle pigments (VAZ) to chlorophyll (C) and the de-epoxidation state of VAZ (D). De-epoxidation state was calculated as (Z+0.5A)/(V+A+Z). All data are means of 3 samples, errors bars statndards deviation. The result obtained for lines RNAi-W1-1 and RNAi-W1-9 are similar only the result of RNAi-W1-1 are therefore shown.

Moreover the seedlings of the RNAi-W1 plants showed in contrast to the wild type bleaching and a reduction of the chlorophyll content at 200 µmol photons m^-2^s^-1^ (Fig. 1B). The reduction was more prominent in case of the RNAi-W1-7 line, having the lowest level of WHIRLY1 protein (Krupinska et al., 2014), as compared to the two other lines.

Analyses of carotenoids showed that in leaves of the RNAi-W1 plants the ratio of VAZ (V=violaxanthin, A=antheraxanthin, Z=zeaxanthin) pool pigments to chlorophylls was enhanced at irradiances of 200 and 350 µmol photons m^-2^s^-1^ (Fig. 1C). The enhanced ratio of VAZ/chlorophyll in the RNAi-W1 plants coincided with a higher de-epoxidation state of the VAZ pool (Fig. 1D) indicating synthesis of zeaxanthin from violaxanthin. In line RNAi-W1-7 with the most extreme knockdown of WHIRLY1, the alterations were more dramatic than in line RNAi-W1-1. At low light, no differences were detected between wild type and RNAi-W1 plants indicating that the alterations in the pigment composition are due to high light stress.

Zeaxanthin is known to have the highest antioxidative capacity of the xanthophylls and might protect thylakoid membrane lipids from oxidation (Havaux et al., 2007). Besides its direct effect as ROS scavenger, zeaxanthin plays an important role in non-photochemical quenching dissipating excess energy as heat and avoiding thereby the production of reactive oxygen species (Li et al., 2009). The enhanced de-epoxidation of the xanthophyll cycle pigments in the RNAi-W1 plants compared to the wild type therefore indicates that their photosynthetic apparatus absorbed more light than required for assimilation of carbon. ROS production by thylakoids from RNAi-W1 or from wild-type seedlings grown at 200 µmol photons m^-2^ s^-1^ was measured by electron paramagnetic spin resonance (EPR). Indirect spin trapping of superoxide/hydrogen peroxide using 4-POBN/ethanol/FeEDTA (Mubarakshina et al., 2010)

Showed that RNAi-W1 thylakoids generated in the light about two times larger signals as wild-type thylakoids (Fig. 2A, B). To investigate whether also singlet oxygen production by thylakoids is enhanced in WHIRLY1 deficient chloroplasts, EPR measurements were performed with the specific spin probe TMPD (Krieger-Liszkay et al., 2015). Using TMPD as spin trap, no difference was observed between the wild type and the transgenic lines (Fig. 2A).

**Figure 2.**
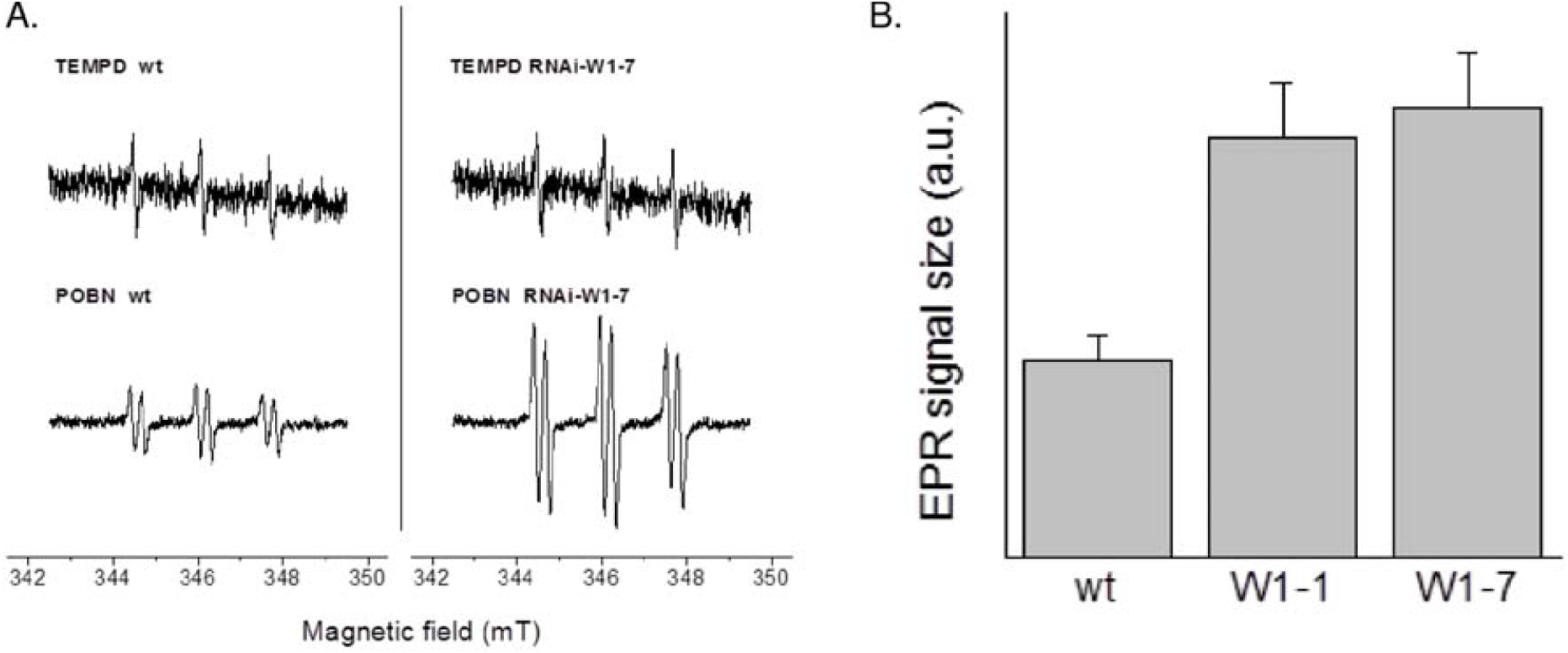
Ros production by thylakoids from waild type and RNAi-W1-1 and RNAi-W1-7 lines. Thylakoids were prpared from seedling grown in continuous light of 200 µ mol photons m^-1^s^-1^. Superoxide /Hydrogen preoxide levels measured by spin trapping EPR using 4-POBN/ErOH/FeEDTA as spintrap and singlet oxygen by the spin probe TMPD-HCl (for experimental details see Krieger – Liszkay et al., 2015) Thylakoids were illuminated for 2 min with res light (500 µmol quanta m^-1^s^-1^) in presence of the spintraps. Left representaticve spectra, right: Siza of the EPR singnals (4-POBN/EtOH/FeEDTA) were normalized to the signal obtained in wild-type thylakoids(mean ± SD, n=6)

Taken together, analyses of pigments as well as ROS measurements revealed that the WHIRLY1 deficient plants experienced more photooxidative stress than the wild type when grown in continuous high light. This indicates that WHIRLY1, in addition to its positive effect on growth, also promotes stress resistance.

Since WHIRLY1 in chloroplasts was shown to bind to RNA as well as to DNA (Melonek et al., 2010) it was obvious to investigate a putative role of WHIRLY1 in controlling the levels of microRNAs which play a central role in the control of plant development as well as in stress responses (Kruszka et al., 2012; Li et al., 2016). For the analysis of microRNAs, primary foliage leaves of wild-type plants and plants of the RNAi-W1-7 line, respectively, grown either at low light (100 µmol photons m^-2^ s^-1^) or at high light (350 µmol photons m^-2^ s^1^), were used. Eight conserved microRNAs reported to be stress responsive in *Arabidopsis thaliana* (Barciszweska et al., 2015) were selected and their levels were determined by Northern blot analyses as well as by RT-qPCR TaqMan MicroRNA assays.

In wild-type plants the levels of most of these microRNAs were enhanced at high light compared to low light conditions (Supplemental Fig. S1). These findings were confirmed by RT-qPCR TaqMan MicroRNA assays, although the changes were not statistically significant in each case (Fig. 3A). While in Northern blot analyses at least several members of a microRNA family were detected (Supplemental Fig. S1), in RT-qPCR TaqMan MicroRNA assays only specific members of a family were measured. Therefore the results of both approaches are not always directly comparable, e.g. in case of miRNA159.

**Figure 3.**
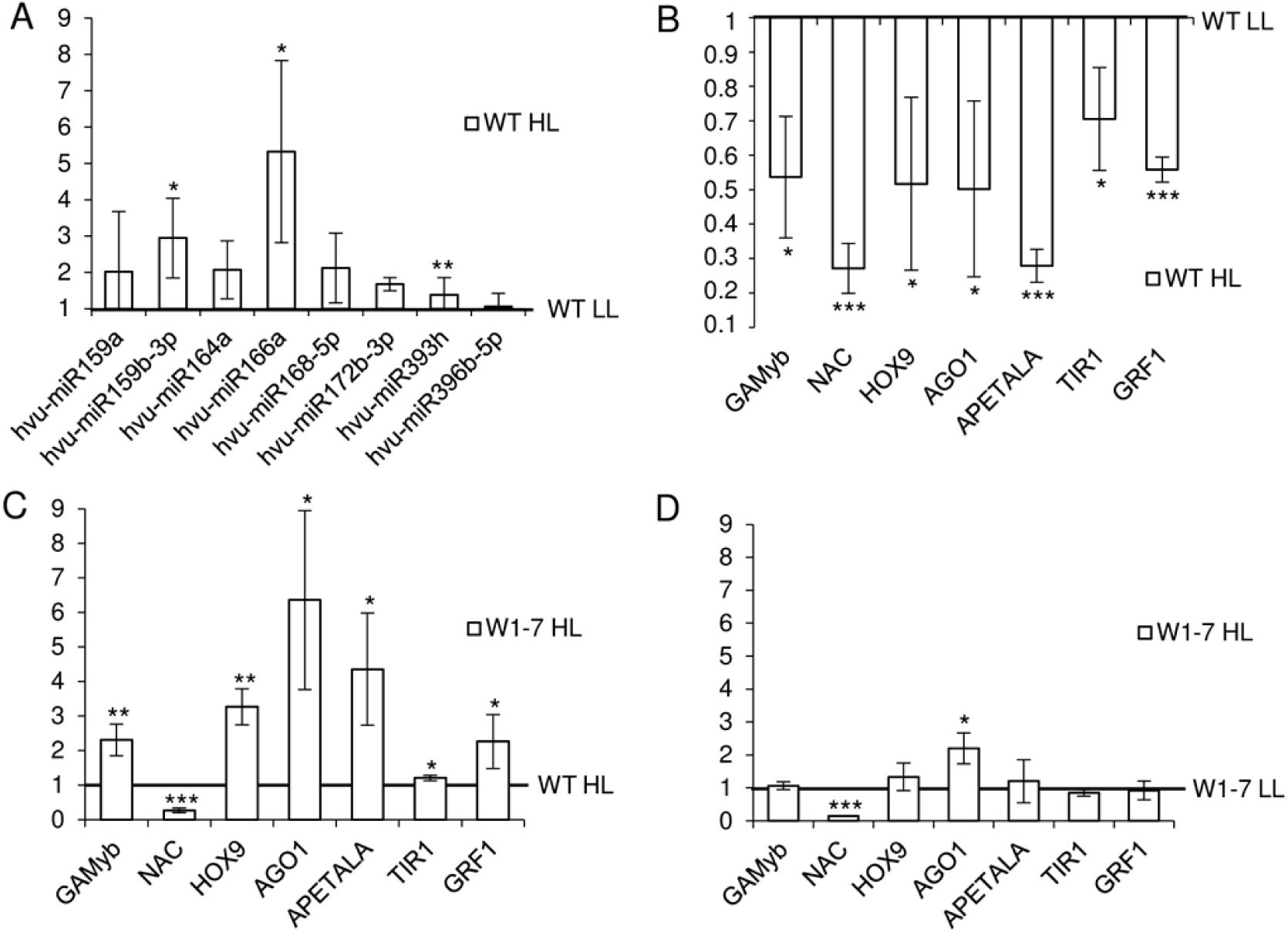
RT–qPCR analysis of microRNAs and target genes expression in wild type (WT) and transgenic RNAi-W1-7 plants exposed to either low(LL) or high light (HL). (A) In WT plants exposed to HL the levels of microRNAs were enhanced. Result are presented as fold change and result for WT plants grows in LL are treated as 1.(B) In the wild type plants HL lead to a downregulation of the levels of traget mRNAs.(C) Levels of most target mRNAs were upregulated in high light treated W1-7 plants exposed to LL and HL are compared. Error bars indicate SD(n=3), and the asterisk indicated a significant different between the sample and control (t test, *P≤0.05, * *P≤0.01, * * *P≤0.001)

For all microRNAs tested in Northern blot hybridization the levels were reduced in leaves of the two RNAi-W1 lines (RNAi-W1-1 and RNAi-W1-7) (Supplemental Fig. S1). These results were confirmed by RT-qPCR TaqMan MicroRNA assays and were independent of the light conditions (Supplemental Fig. S2A, B).

Some mRNA targets for the tested microRNAs are known in several plant species including barley (Supplemental Table 1). Additional missing target mRNAs in barley were identified using the psRNA-Target software (Dai and Zhao, 2011; <u>http://plantgrn.noble.org/psRNATarget/</u>) (Supplemental Material S2). MicroRNAs hvu-miR159a and hvu-miR159b-3p potentially target *GAMyb* mRNA, and microRNAs named hvu-miR164a, hvu-miR172b-3p, hvu-miR393h and hvu-miR396b-5p target *NAC*, *APETALA*, *TIR1* and *GRF1* mRNAs, respectively. Barley *HOX9* and *AGO1* mRNAs have been shown experimentally to be targets of microRNA166a and microRNA168-5p, respectively (Kruszka et al., 2014, Pacak et al., 2016).

The effects of the selected miRNAs on the levels of targeted mRNAs were tested by qRT-PCR. In primary foliage leaves of wild-type plants grown in high light, the upregulation of microRNAs coincided with a downregulation of targeted mRNAs (Fig. 3B). In contrast, in the RNAi-W1 plants grown in high light most target gene mRNA levels are enhanced compared to the wild type (Fig. 3C) whereas at low light the levels of target gene mRNAs are similar between the wild type and RNAi-W1-7 plants (Supplemental Fig. S3).

The results indicate that high light induced signals from chloroplasts stimulate a WHIRLY1 dependent downregulation of the level of mRNAs targeted by the tested microRNAs being upregulated in the wild type. In contrast to the wild type, plants of the RNAi-W1-7 line did neither show a light-induced increase in microRNAs nor a decrease in the mRNA levels of their target genes. This indicates that the WHIRLY1 deficient plants can’t respond to stress and thereby suffer from a higher ROS production.

In the transgenic plants grown either in low light or in high light, the levels of targeted mRNAs did not show essential differences (Fig. 3D). The only exception is NAC transcription factor mRNA (GenBank: AK356223.1) that is downregulated in W1-7 high and low light grown plants despite the low level of its potential cognate microRNA164a. The reason for this result remains unclear. NAC transcription factors comprise one of the largest gene families and are involved in the regulation of plant development, senescence and response to various stresses. Their activities can be regulated at different levels (transcription efficiency, alternative splicing, posttranslational regulation) that possibly might affect the final level of NAC mRNAs (Shao et al., 2015).

WHIRLY1 has been proposed to move from the chloroplast to the nucleus in response to environmental cues such as high light intensity (Foyer et al., 2014). In this study it has been demonstrated that the repertoire of the plants’ responses towards high light involves a WHIRLY1 dependent increase in the levels of diverse nuclear microRNAs. As WHIRLY1 can bind to RNA it might be a general factor influencing the biogenesis and/or stability of microRNAs. The observed phenomenon might be caused either by direct binding of WHIRLY1 to the nuclear microRNAs and/or its architectural impact on nuclear chromatin as observed in chloroplasts (Krupinska et al., 2014). To elucidate the specific role of WHIRLY1 in the regulation of the levels of microRNAs and targeted mRNAs during retrograde signaling further detailed studies are required.

## Acknowledgements

We would like to thank Artur Jarmolowski (AMU Poznan, Poland) for fruitful discussions. Susanne Braun an Jens Hermann (CAU Kiel, Germany) are thanked for technical assistance. We thank the German Research Foundation (DFG: Kr1350/7, Kr1350/9), National Science Centre Poland (NSC UMO-2016/23/B/NZ9/00862), and the KNOW RNA Research Centre in Poznan 01/KNOW2/2014 for financial support.

## SUPPLEMENTAL DATA

**Supplemental Figure S1**. Northern blot analysis of microRNA levels in low light (LL) and high light (HL) in wild type (WT) and WHIRLY1 deficient barley plants (RNAi-W1-1 and RNAi-W1-7).

**Supplemental Figure S2**. RT-qPCR analysis of microRNAs in wild type (WT) and transgenic RNAi -W1-7 plants exposed to either low (LL) or high light (HL).

**Supplemental Figure S3**. RT-qPCR analysis of target genes expression in wild type (WT) and transgenic RNAi -W1-7 plants exposed to either low light (LL).

**Supplemental Table 1**. List of microRNAs, their sequences, NCBI GEO accession numbers of barley Next Generation Sequencing results and references.

**Supplemental Table 2**. List of microRNA sequences, TaqMan™ MicroRNA assays and Northern probes used in the study.

**Supplemental Table 3**. Primer sequences used in the RT-qPCR of target mRNA levels.

**Supplemental Material and Methods S1**. A supplemental “Materials and Methods” section.

**Supplemental Material S2.** psRNA-Target analysis results.

